# RNA sequence to structure analysis from comprehensive pairwise mutagenesis of multiple self-cleaving ribozymes

**DOI:** 10.1101/2022.05.17.492349

**Authors:** Jessica M. Roberts, James D. Beck, Tanner B. Pollock, Devin P. Bendixsen, Eric J. Hayden

## Abstract

Self-cleaving ribozymes are RNA molecules that catalyze the cleavage of their own phosphodiester backbones. These ribozymes are found in all domains of life and are also a tool for biotechnical and synthetic biology applications. Self-cleaving ribozymes are also an important model of sequence to function relationships for RNA because their small size simplifies synthesis of genetic variants and self-cleaving activity is an accessible readout of the functional consequence of the mutation. Here we used a high-throughput experimental approach to determine the relative activity for every possible single and double mutant of five self-cleaving ribozymes. From this data, we comprehensively identified non-additive effects between pairs of mutations (epistasis) for all five ribozymes. We analyzed how changes in activity and trends in epistasis map to the ribozyme structures. The variety of structures studied provided opportunities to observe several examples of common structural elements, and the data was collected under identical experimental conditions to enable direct comparison. Heat-map based visualization of the data revealed patterns indicating structural features of the ribozymes including paired regions, unpaired loops, non-canonical structures and tertiary structural contacts. The data also revealed signatures of functionally critical nucleotides involved in catalysis. The results demonstrate that the data sets provide structural information similar to chemical or enzymatic probing experiments, but with additional quantitative functional information. The large-scale data sets can be used for models predicting structure and function and for efforts to engineer self-cleaving ribozymes.

## INTRODUCTION

Challenges with predicting the functional effects of changing an RNA sequence continues to limit the study and design of RNA molecules. Recently, machine learning approaches have made considerable advancements in predicting an RNA structure from a sequence. However, these approaches rely heavily on crystal structures of RNA molecules and sequence conservation of homologs, both of which are limited for RNA molecules compared to proteins (Calonaci et al., 2020; Townshend et al., 2021). In addition, describing an RNA molecule as a single structure can be inaccurate, and regulatory elements such as riboswitches demonstrate the importance of an ensemble of structures for an RNA function. It is unclear that predictions based on individual structures alone will be able to predict functional effects of mutations with the precision needed for many biotechnical and synthetic biology applications, or to predict disease-associated mutations in RNA molecules (Halvorsen et al., 2010). This suggests that new experimental data types might be important for understanding, designing, and manipulating the transcriptome.

Self-cleaving ribozymes provide a useful model to study sequence-structure-function relationships in RNA molecules. Self-cleaving ribozymes are catalytic RNA molecules that cleave their own phosphodiester backbone. They were first discovered in viruses and viroids, but numerous families of self-cleaving ribozymes have since been discovered in all domains of life (Prody et al., 1986). The CPEB3 ribozyme, for example, was discovered in the human genome and found to be highly conserved in mammals (Bendixsen et al., 2021, p. 3; Salehi-Ashtiani et al., 2006). Other self-cleaving ribozymes, such as the hammerhead and twister ribozymes, are found broadly distributed across eukaryotic and prokaryotic genomes (Perreault et al., 2011;

Roth et al., 2014). The biological roles of ribozymes in different genomes and different genetic contexts remain an active area of investigation (Jimenez et al., 2015). In addition to being widespread across the tree of life, self-cleaving ribozymes have also been used for several bioengineering applications (Liang et al., 2011; Peng et al., 2021; Wei and Smolke, 2015; Zhong et al., 2016). For example, self-cleaving ribozymes are being combined with aptamers to develop synthetic gene regulatory devices, which have biotechnical and biomedical applications where ligand dependent control of gene expression is desired (Kobori et al., 2017, 2015; Stifel et al., 2019; Townshend et al., 2015).

The testing of mutational effects in ribozyme sequences has been accelerated by high-throughput experimental approaches. Most self-cleaving ribozymes are fairly small (<200 nt) and genetic variants can be made by chemical synthesis of a single DNA oligonucleotide that is then used as a template for in vitro transcription. The self-cleavage activity of the ribozyme requires a precise three-dimensional structure, and therefore activity can be used as a sensitive indirect readout of native structure. Mutations that disrupt the native structure are detected as reduced activity compared to the unmutated “wild-type” ribozyme. Several methods have been developed to enable the detection of ribozyme function by high-throughput sequencing of biochemical reactions (Bendixsen et al., 2019; Hayden, 2016; Kobori and Yokobayashi, 2016; Shen et al., 2021). For self-cleaving ribozymes, each read from the data reports both the mutations and whether or not that molecule was reacted (cleaved) or unreacted (uncleaved). Therefore, high-throughput sequencing allows numerous genetic variants to be pooled together and still observed hundreds to thousands of times in the data. This provides confidence in the fraction cleaved for each genetic variant in a given experiment, and genetic variants are compared to determine relative activity. Importantly, the data is internally controlled because both reacted and unreacted molecules are observed, which controls for differences in their abundance due to synthesis steps (chemical DNA synthesis, transcription, reverse-transcription, PCR).

A common approach to confirm structural interactions in RNA and proteins is through analysis of pairs of mutations (Dutheil et al., 2010; Olson et al., 2014). In this context, it can be useful to calculate pairwise epistasis, which measures deviations in the mutational effects of double mutants relative to the effects of each individual mutation (assuming an additive model of mutational effects). For example, in the case of a base-pair, each single mutation would disrupt the base-pairing interaction, destabilizing the catalytically active RNA structure and reducing activity. However, if two mutants together restore a base-pair, the relative activity of the double mutant would have much higher activity than expected from the additive effects of the individual mutations (positive epistasis). In contrast to paired nucleotides, double mutants at non-paired nucleotides tend to have a more reduced activity than expected from each individual mutation (negative epistasis) (Bendixsen et al., 2017; Li et al., 2016). In the case of two mutations that create a different base pair (i.e. G-C to A-U), it is known that the stacking with neighboring base pairs is also structurally important, and some base pair substitutions will not be equivalent in a given structural context. This creates a range of possible epistatic effects even for two mutations at paired nucleotide positions. In addition, some non-canonical base interactions within tertiary contacts may also show epistasis even when they do not involve Watson-Crick or GU wobble base pairing interactions. Nevertheless, the propensity for positive epistasis between physically interacting nucleotides suggests that a comprehensive evaluation of pairwise mutational effects should contain considerable structural information.

Here, we report comprehensive analysis of mutational effects for all single and double mutants for five different self-cleaving ribozymes. Relative activity effects of all single and double mutations were determined by high-throughput sequencing of co-transcriptional self-cleavage reactions, and this data was used to calculate epistasis between pairs of mutations. The ribozymes studied include a mammalian CPEB3 ribozyme, a Hepatitis Delta Virus (HDV) ribozyme, a twister ribozyme from *Oryza sativa*, a hairpin ribozyme derived from the satellite RNA from tobacco ringspot virus, and a hammerhead ribozyme (Bendixsen et al., 2021; Burke and Greathouse, 2005; Chadalavada et al., 2007; Liu et al., 2014; Müller et al., 2012). For each reference ribozyme, a single DNA oligo template library was synthesized with 97% wild-type nucleotides at each position, and 1% of each of the three other nucleotides. This mutagenesis strategy was expected to produce all possible single and double mutants, as well as a random sampling of combinations of three or more mutations. The mutagenized templates were transcribed in vitro, all under identical conditions, where active ribozymes had the opportunity to self-cleave co-transcriptionally. All ribozyme constructs studied cleave near the 5’-end of the RNA, and a template switching reverse transcription protocol was used to append a common primer binding site to both cleaved and uncleaved molecules. Subsequently, low cycle PCR was used to add indexed Illumina adapters for high-throughput sequencing. Each mutagenized ribozyme template was transcribed separately and in triplicate, and amplified with unique indexes so that all replicates could be pooled and sequenced together on an Illumina sequencer. The sequencing data was then used to count the number of times each unique sequence was observed as cleaved or uncleaved, and this data was used to calculate the fraction cleaved. The fraction cleaved of single and double mutants was normalized to the unmutated reference sequence to determine relative activity. The relative activity values of the single and double mutants were used to calculate all possible pairwise epistatic interactions in all five ribozymes. We mapped epistasis values to each ribozyme structure to evaluate correlations between structural elements and patterns of pairwise epistasis values. The results indicated that structural features of the ribozymes are revealed in the data, suggesting that these data sets will be useful for developing models for predicting sequence-structure-function relationships in RNA molecules.

## RESULTS AND DISCUSSION

### Evaluation of read depth and mutational coverage

The accuracy of our relative activity measurements depends on the number of reads we observe that map to each unique ribozyme sequence (read depth). Each reference ribozyme has a different nucleotide length resulting in different numbers of possible single and double mutants. In addition, the pooling of experimental replicates for sequencing does not result in equal mixtures of each replicate. In order to determine read depth, we mapped reads to the reference sequences and counted the number of reads that matched each ribozyme, while allowing for 1 or 2 mutations. We observed every single and double mutant for all ribozymes in each replicate, indicating 100% coverage of these mutant classes for all of our data sets. The distributions of observations for each single and double mutant of each ribozyme are shown in Supplementary Figure 1. The HDV data showed the lowest depth, possibly because it is a larger ribozyme (87 nt), and fewer reads mapped to the single and double mutants (Table 1). Nevertheless, from this analysis we conclude that the data contains complete coverage of all single and double mutants and ample read depth for all five ribozymes.

**Table 1:** Summary of the lengths of each self-cleaving ribozyme used in this study, and the number of single and double mutants whose cleavage activity was analyzed.

| **Ribozyme Name** | **Ribozyme Length** | **Possible Single Mutants** | **Possible Double Mutants** | **Total Mapped Reads** | **Overall Fraction Cleaved** |
| --- | --- | --- | --- | --- | --- |
| **CPEB3** | **69** | **207** | **21,114** | **9,238,603** | **0.68** |
| **HDV** | **87** | **261** | **33,669** | **3,316,380** | **0.60** |
| **Twister** | **48** | **144** | **10,152** | **7,762,863** | **0.31** |
| **Hairpin** | **71** | **213** | **22,365** | **5,067,216** | **0.23** |
| **Hammerhead** | **45** | **135** | **8,910** | **8,054,498** | **0.19** |

### Epistatic effects in paired nucleotide positions show stability-dependent signatures

In order to evaluate how the effects of mutations mapped to the ribozyme structures, we plotted the relative activity values as heat maps (Figures 1-5). We then used this data to calculate epistasis between pairs of mutations. We first inspected nucleotide positions known to be involved in base-paired regions of the secondary structure of each ribozyme. In this heatmap layout, many paired regions showed an anti-diagonal line of high activity double mutant variants with strong positive epistasis (Figures 1-5, insets). In addition, pairs of mutations off the anti-diagonal tended to show negative or non-positive epistasis. Pseudoknot elements that involve Watson-Crick base pairs also showed this pattern, including the single base pair T1 element in CPEB3 (Figure 1) and the two base pair T1 element in HDV (Figure 2). The layout of mutations in the heatmap places paired nucleotide positions along the anti-diagonal and compensatory double mutants that change one Watson-Crick base pair to another are found on this anti-diagonal. Individual mutations that break a base pair will often reduce ribozyme activity, but the activity can be restored by a second compensatory mutation resulting in positive epistasis. In contrast, double mutants off-diagonal usually disrupt two base pairs (unless they result in a GU wobble base pair). It is expected that breaking two base pairs in the same paired region would be more deleterious to ribozyme activity than breaking one base pair, but it appears that two non-compensatory mutations in the same paired region are more deleterious than expected from an additive assumption, and frequently create negative epistasis off-diagonal within paired regions.

**Figure 1:**
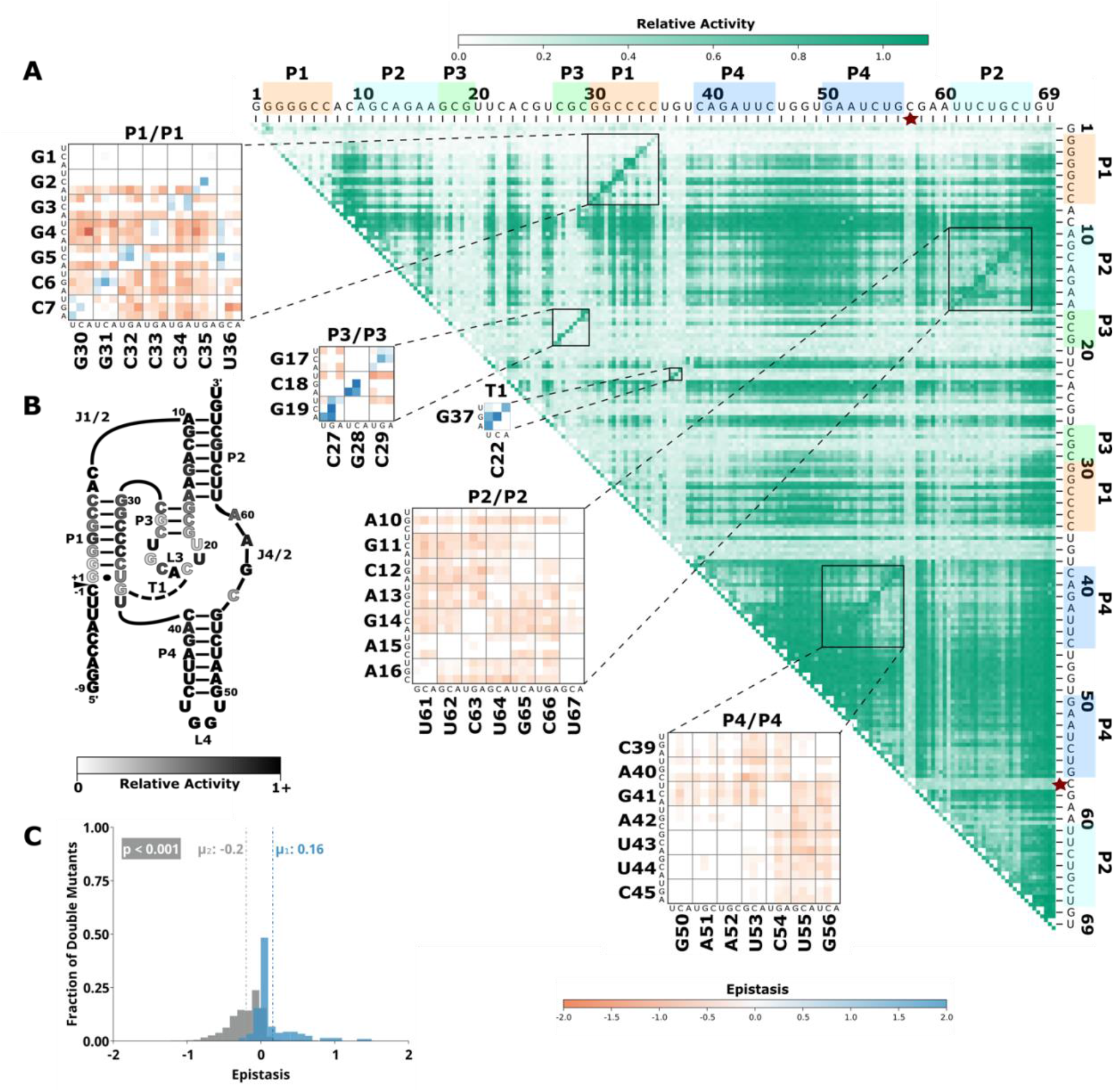
Effects of mutations and pairwise epistasis in a CPEB3 ribozyme. A) Relative activity heatmap depicting all possible pairwise effects of mutations on the cleavage activity of a mammalian CPEB3 ribozyme. Base-paired regions P1, P2, P3, P4, and T1 are highlighted and color coordinated along the axes, and surrounded by black squares within the heatmap. Pairwise epistasis interactions observed for each paired regions are each shown as expanded insets for easy identification of the specific epistatic effects measured for each pair of mutations. Instances of positive epistasis are shaded blue, and negative epistasis is shaded red, with higher color intensity indicating a greater magnitude of epistasis. Catalytic residues are indicated by stars along the axes. B) Secondary structure of the CPEB3 ribozyme used in this study. Each nucleotide is shaded to indicate the average relative cleavage activity of all single mutations at that position. C) Histogram showing the distributions of epistasis in the paired regions of CPEB3. The distribution for double mutants within a paired region that are not involved in a base-pair is shown in grey, and the distribution for nucleotides involved in a base-pair is shown in blue.

**Figure 2:**
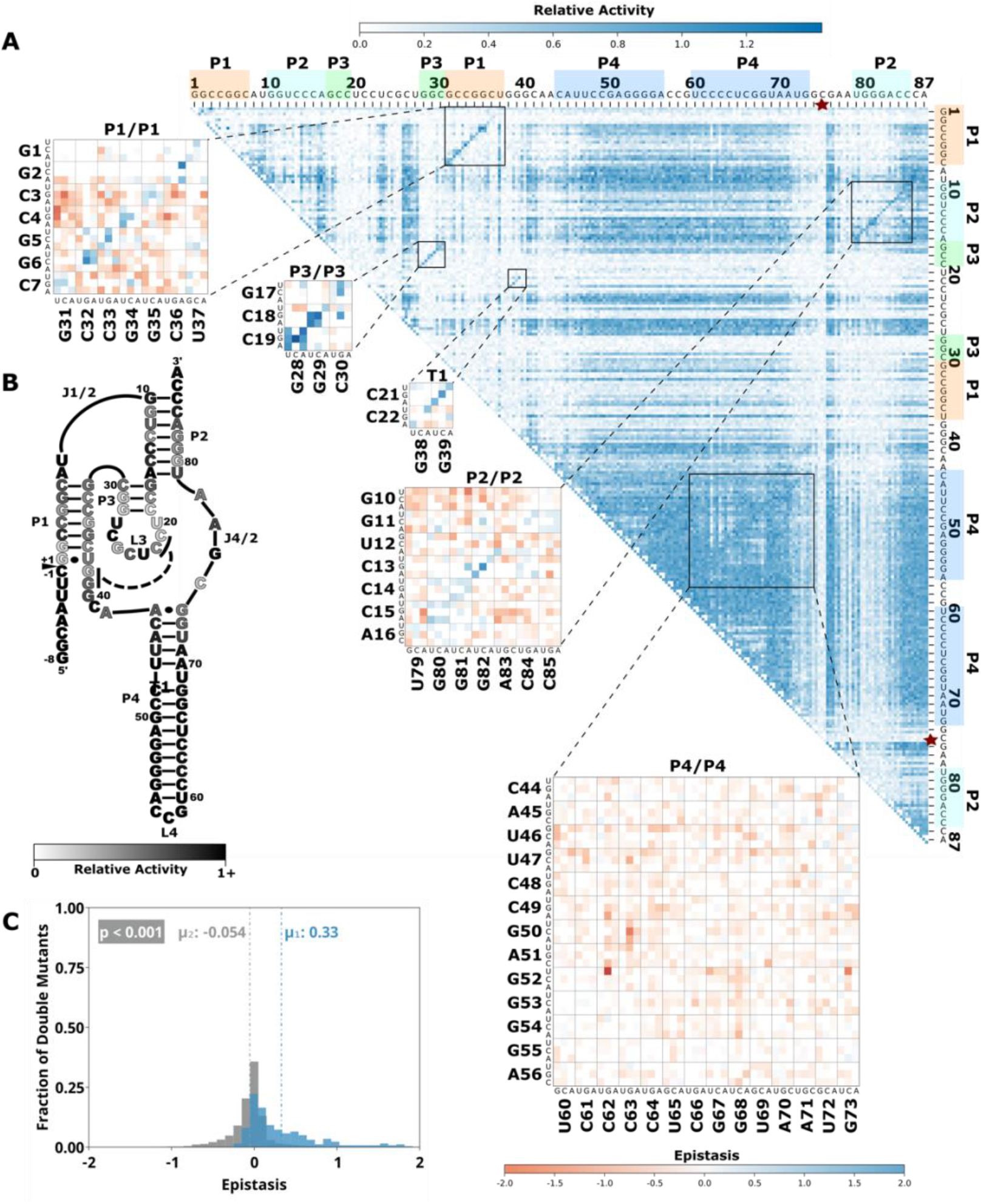
Comprehensive pairwise epistasis landscape for a HDV self-cleaving ribozyme. A) Relative activity heatmap depicting all possible pairwise effects of mutations on the cleavage activity of an HDV ribozyme. Base-paired regions P1, P2, P3, P4, and T1 are highlighted and color coordinated along the axes, and surrounded by black squares within the heatmap. Pairwise epistasis interactions observed for each paired regions are each shown as expanded insets for easy identification of the specific epistatic effects measured for each pair of mutations. Instances of positive epistasis are shaded blue, and negative epistasis is shaded red, with higher color intensity indicating a greater magnitude of epistasis. Catalytic residues are indicated by stars along the axes. B) Secondary structure of the HDV ribozyme used in this study. Each nucleotide is shaded to indicate the average relative cleavage activity of all single mutations at that position. C) Histogram showing the distributions of epistasis in the paired regions of HDV. The distribution for double mutants within a paired region that are not involved in a base-pair is shown in grey, and the distribution for nucleotides involved in a base-pair is shown in blue.

**Figure 3:**
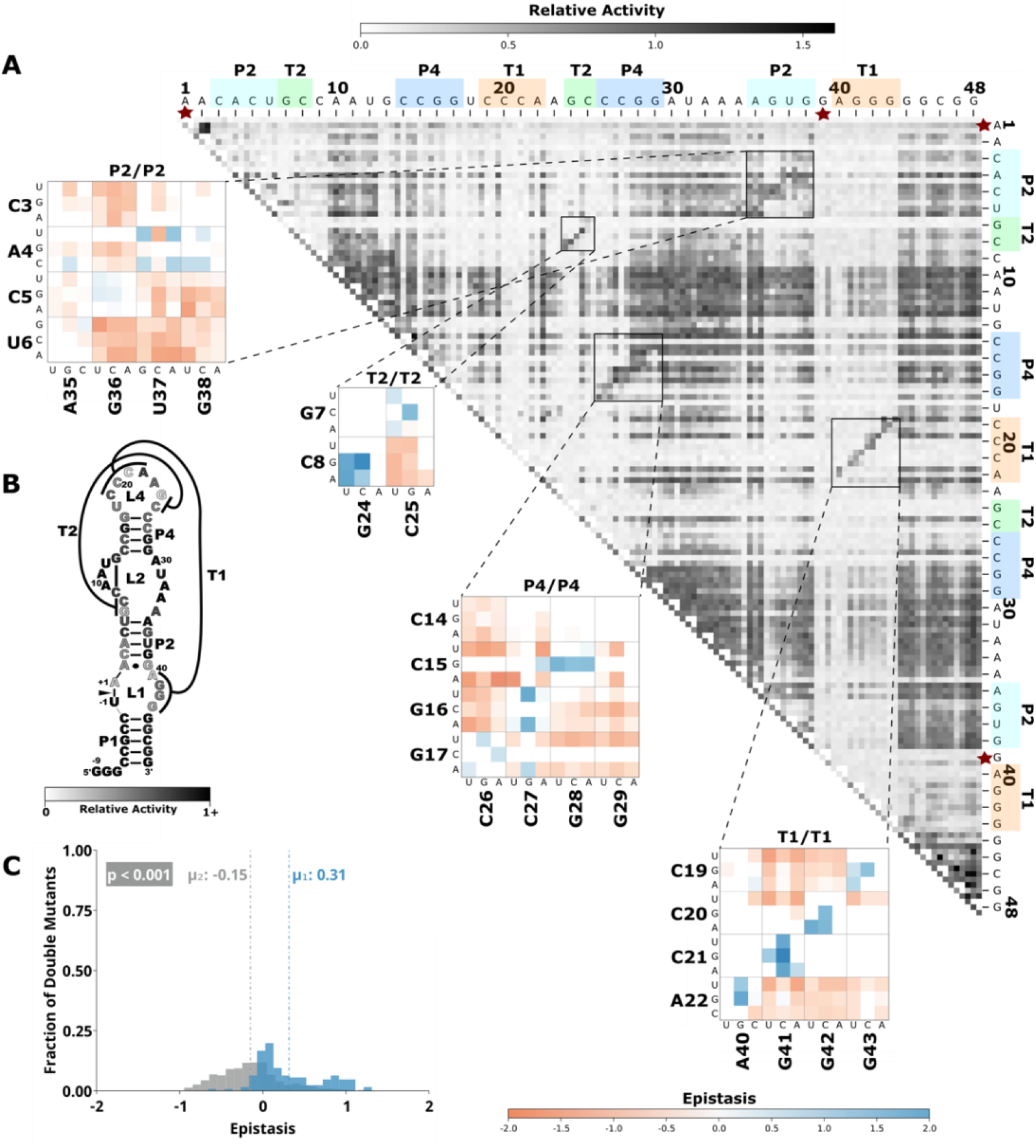
Comprehensive pairwise epistasis landscape for a twister self-cleaving ribozyme. A) Relative activity heatmap depicting all possible pairwise effects of mutations on the cleavage activity of a twister ribozyme. Base-paired regions P2, P4, T1, and T2 are highlighted and color coordinated along the axes, and surrounded by black squares within the heatmap. Pairwise epistasis interactions observed for each paired region are each shown as expanded insets for easy identification of the specific epistatic effects measured for each pair of mutations. Instances of positive epistasis are shaded blue, and negative epistasis is shaded red, with higher color intensity indicating a greater magnitude of epistasis. Catalytic residues are indicated by stars along the axes. B) Secondary structure of the twister ribozyme used in this study. Each nucleotide is shaded to indicate the average relative cleavage activity of all single mutations at that position. C) Histogram showing the distributions of epistasis in the paired regions of twister. The distribution for double mutants within a paired region that are not involved in a base-pair is shown in grey, and the distribution for nucleotides involved in a base-pair is shown in blue.

**Figure 4:**
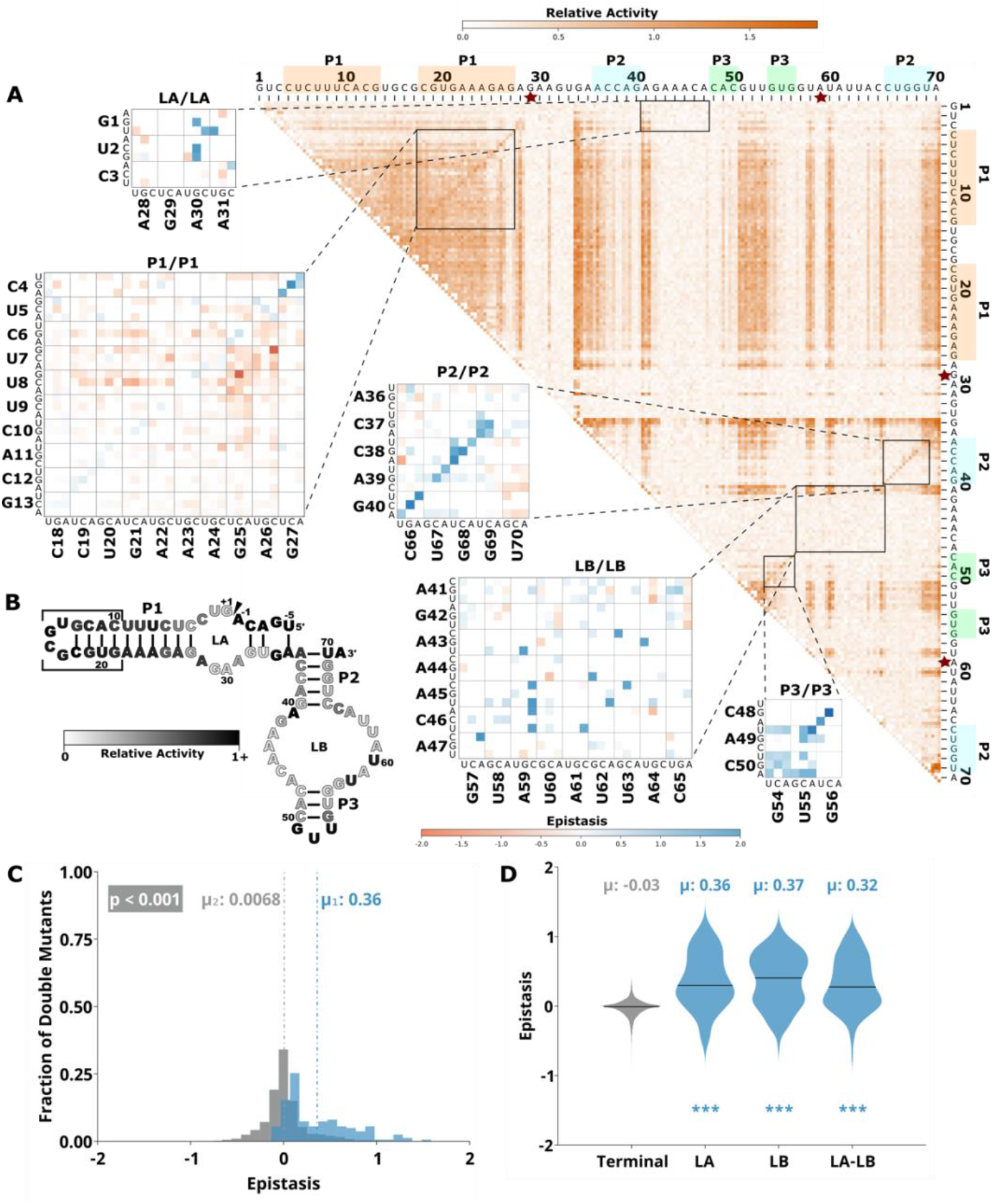
Comprehensive pairwise epistasis landscape for a hairpin self-cleaving ribozyme. A) Relative activity heatmap depicting all possible pairwise effects of mutations on the cleavage activity of a hairpin ribozyme. Base-paired regions P1, P2, and P3 are highlighted and color coordinated along the axes, and surrounded by black squares within the heatmap. Pairwise epistasis interactions observed for each paired region are each shown as expanded insets for easy identification of the specific epistatic effects measured for each pair of mutations. Instances of positive epistasis are shaded blue, and negative epistasis is shaded red, with higher color intensity indicating a greater magnitude of epistasis. Catalytic residues are indicated by stars along the axes. B) Secondary structure of the hairpin ribozyme used in this study. Each nucleotide is shaded to indicate the average relative cleavage activity of all single mutations at that position. C) Histogram showing the distributions of epistasis in the paired regions of hairpin. The distribution for double mutants within a paired region that are not involved in a base-pair is shown in grey, and the distribution for nucleotides involved in a base-pair is shown in blue.

**Figure 5:**
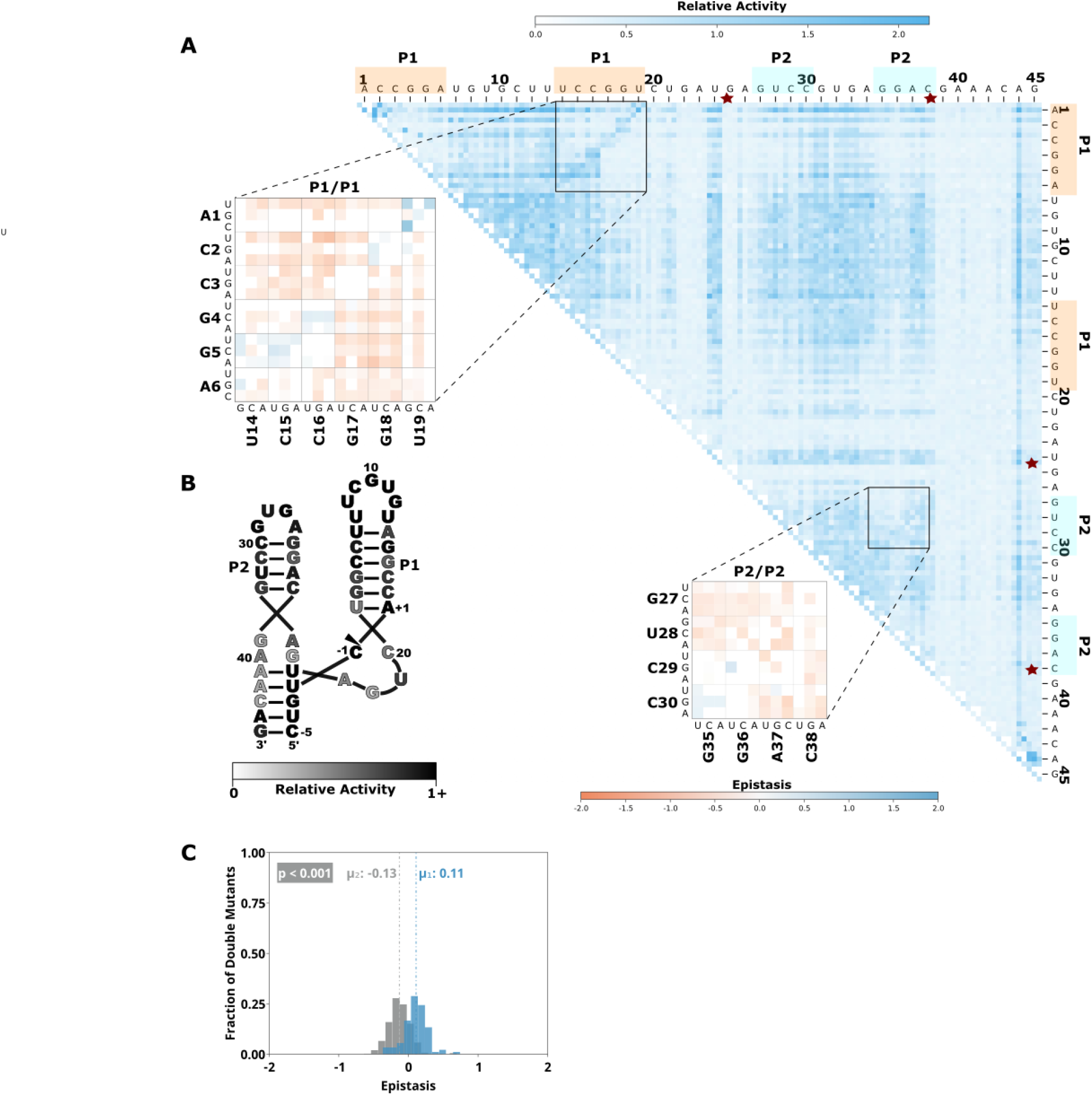
Comprehensive pairwise epistasis landscape for a hammerhead self-cleaving ribozyme. A) Relative activity heatmap depicting all possible pairwise effects of mutations on the cleavage activity of a hammerhead ribozyme. Base-paired regions P1, and P2 are highlighted and color coordinated along the axes, and surrounded by black squares within the heatmap. Pairwise epistasis interactions observed for each paired region are each shown as expanded insets for easy identification of the specific epistatic effects measured for each pair of mutations. Instances of positive epistasis are shaded blue, and negative epistasis is shaded red, with higher color intensity indicating a greater magnitude of epistasis. Catalytic residues are indicated by stars along the axes. B) Secondary structure of the hammerhead ribozyme used in this study. Each nucleotide is shaded to indicate the average relative cleavage activity of all single mutations at that position. C) Histogram showing the distributions of epistasis in the paired regions of hammerhead. The distribution for double mutants within a paired region that are not involved in a base-pair is shown in grey, and the distribution for nucleotides involved in a base-pair is shown in blue.

To quantify the observed difference in epistasis between nucleotide positions that form a base pair and two that do not, we plotted the distribution of epistasis values for double mutants on and off the anti-diagonal within the paired regions of each ribozyme. Statistical analysis indicated that the distributions were significantly different (p<0.001, Mann-Whitney U-test), and the epistasis values between paired nucleotide positions (on-diagonal) were consistently more positive than two mutations in positions that are not directly base paired (off-diagonal). This analysis was consistent for every individual paired region in each ribozyme (Figures 1-5, panel C). This pattern of epistasis in paired regions demonstrates the utility of comprehensive double-mutant activity data for identifying base paired regions in RNA structures.

It is interesting to note that the magnitude of the difference in the distributions of epistasis values for double mutants at paired and non-paired positions was different for different paired regions (Supplementary Figure 2). Specifically, short paired elements with fewer base pairs seemed to show large differences in the distributions of epistatic effects for paired and unpaired positions, while longer paired elements showed small differences in these distributions. For example, the short P3 (3 bp) in CPEB3 and HDV, and T1 (4 bp) in the twister ribozyme showed very large differences between the distributions of epistasis values at paired versus non-paired positions. These small regions are highly sensitive to mutations, and most pairs of mutations within this region result in almost no detectable activity except when they create a different Watson-Crick base pair (Figures 1-5). These structural elements have positive epistasis along the anti-diagonal, and negative epistasis off diagonal, resulting in large differences between the distributions of epistasis (Supplementary Figure 2). In contrast, the P4 stem in HDV has the most base pairs of any paired region in this data set (14), and losing one of these base pairs was not deleterious to riboyzme activity in our experiments (Figure 2). Because the single mutations had little effects on the self-cleavage activity, a compensatory mutation restoring a base pair did not result in positive epistasis (Figure 2). Futher, only weak negative epistasis is observed off-diagonal indicating that the loss of two base pairs in P4 was somewhat tollerated compared to shorter paired regions. The distributions for epistatis for paired and unpaired positions in P4 of HDV show only a small difference (Supplementary Figure 2). Together, the differences between epistasis in short and long base paired regions suggests that the thermodynamic stability of each paired region is important for the observed activity differences contributing to epistasis, which might ultimately affect the utility of this data for identifying paired regions in RNA structures.

In order to quantify the influence of thermodynamic stability on epistasis in different paired regions, we calculated the minimum free energy for each paired region and compared mutational effects. We split each paired region into two separate RNA sequences that contained only the base paired nucleotides and used nearest neighbor rules to calculate the minimum free energy of their interaction (NUPACK). This approach neglects thermodynamic contributions from terminal loops, but allowed for a consistent approach to compare internal and terminal paired regions. We found a significant negative correlation between the median deleterious effects of single mutations and the minimum free energy of the paired regions (Supplementary Figure 3). This analysis indicates that more stable structural elements may be harder to identify from epistatic effects. However, it is possible that more stable elements would show stronger epistasis under different experimental conditions, such as different temperatures or magnesium concentrations (Peri et al., 2022).

### Catalytic residues do not have any high-activity mutants, and do not exhibit epistasis

Self-cleaving ribozymes often utilize a concerted acid base catalysis mechanism where specific nucleobases act as proton donors (acid) or acceptors (base) (Jimenez et al., 2015), and mutations at these positions abolish activity. Analyzing the effects of individual mutations will not distinguish catalytic nucleotides from structurally important nucleotides. Comprehensive pairwise mutations, on the other hand, can potentially distinguish between structurally important nucleotides involved in paired regions that show positive epistasis from compensatory effects. The catalytic cytosines of the CPEB3 (C57) and HDV (C75) act as proton donors due to perturbed pKa values (Nakano, 2000; Skilandat et al., 2016). For the twister ribozyme (Figure 3) the guanosine at position G39 acts as a general base, and the adenosine at position A1 acts as a general acid (Wilson et al., 2016). The catalytic nucleotides for the Hammerhead ribozyme (Figure 5) are the Guanosines located at positions G25 and G39 (Scott et al., 2013). The hairpin ribozyme (Figure 4) contains catalytic nucleotides at positions G29 and A59 (Wilson, 2006). In the relative activity heat maps, the columns and rows associated with these nucleotides result in low activity values (Figures 1-5, Supplementary Figure 4). It is important to note that because there is complete coverage of all double mutants in this data set, we can be certain that there are no possible compensatory mutations. These results show how catalytic residues can be identified in the comprehensive pairwise mutagenesis data.

### Unpaired nucleotides show tertiary structure dependent mutational effects

Mutations to nucleotides found in terminal loops that are not involved in tertiary structure elements showed high relative activity for most single and double mutants, and essentially no epistasis. This is not surprising if these loops reside on the periphery of the ribozyme and are not involved in structural contacts with other nucleotides. This is the case for L4 of the CPEB3 and HDV ribozymes (Figure 1, Figure 2), and L1 and L3 of the hairpin ribozyme (Figure 4). Two mutations within these loops do not reduce activity, and mutations in these loops do not rescue other deleterious mutations such as those that break a base pair (Figures 1, 2, and 4).

The internal loops (LA and LB) of the hairpin ribozyme are structurally important (Figure 4). Interactions between nucleotides within LB include six non-Watson-Crick base pairing interactions that are important for the formation of an active ribozyme structure (Fedor, 2000). Several non-canonical base-base and sugar-base hydrogen bonds between nucleotides within LA are also important for the formation of the active site (Fedor, 2000; Wilson, 2006). Docking between LA and LB is necessary for the formation of a catalytically active ribozyme and is facilitated by a Watson-Crick base pair between G1 and C46 in the version of the ribozyme used here (Rupert and Ferré-D’Amaré, 2001). In contrast to terminal loop regions, most single mutations within LA and LB resulted in low self-cleavage activity in our data (Figure 4). In addition, the double mutants within and between loop A and loop B show several instances of strong positive epistasis (Figure 4, Insets), and the distributions of epistasis within and between these loops are significantly different than the terminal loops that are not structurally important (Figure 4D). This positive epistasis indicates that many of the important structural contacts can be facilitated by other specific pairs of nucleotides. For example, the double mutant G1C and C46G shows strong epistasis suggesting that swapping a C-G base pair for the G-C base pair can restore activity by facilitating docking between the two loops. Several double mutants at positions that form non-canonical interactions in LB show positive epistasis. For example, mutation A41G shows positive epistasis when the interacting nucleotide C65 is mutated to a G or U. The non-canonical base pair G42:A64 shows positive epistasis for the mutations G42U A64G. The non-canonical A45:A59 interaction shows positive epistasis for several pairs of mutations (A45U A59C, A45C A59C, A45G A59U). Finally, the non-canonical base pair A47:G57 in LB, and C3:A28 in LA, both show positive epistasis for double mutants that result in an AU base pair. This analysis indicates that important structural contacts can be achieved with several different nucleotide combinations. The difference between terminal loops and loops with structural importance highlights how activity-based data can help identify non-canonical structures that are challenging to predict computationally, and that might be difficult to identify by other common approaches, such as chemical probing experiments (Walter et al., 2000).

Another example of structurally important unpaired regions can be found in the CUGA uridine turn (U-turn) motif in the hammerhead ribozyme (Figure 5). This CUGA turn forms the catalytic pocket and positions a catalytic cytosine (−1C) at the cleavage site (Doudna, 1995). A crystal structure of the sTRSV ribozyme showed a base pair between the nucleotides corresponding to C20 and G25 in the ribozyme construct used for our experiments (Chi et al., 2008). These two nucleotides showed strong positive epistasis for the mutations C20G and G25C, which substitutes a G:C base pair for the original C:G base pair. All other single and double mutants in this region showed low activity, and no instances of strong positive epistasis within or between this motif (Figure 5). The low activity resulting from mutations in this region confirms the functional importance of this motif, and indicates that this motif cannot be easily formed or rescued by sequences with up to two mutational differences, except for the G:C base pair swap.

Tertiary interactions between loops in the hammerhead ribozyme provide another example of structurally important loop regions. Type III hammerhead ribozymes, like the one used in this study, contain tertiary interactions between nucleotides in the loops of P1 and P2 that are implicated in structural organization of the catalytic core. A crystal structure of this loop-loop interaction showed a network of interhelical non-canonical base pairs and stacks, with several nucleobases in stem-loop I interacting with more than one nucleobase in stem-loop II (Chi et al., 2008; Martick and Scott, 2006). However, there are numerous different loop sequences in naturally occurring hammerhead ribozymes indicating that this loop-loop interaction can be formed by a variety of different sequences (Burke and Greathouse, 2005; Perreault et al., 2011). We therefore anticipated that the we would observe a significant level of positive epistasis between these two loops for double mutations that were capable of maintaining these tertiary interactions. Surprisingly, however, we found that most individual and double mutations do not reduce activity (Figure 5), and double mutants do not show positive epistasis (Supplementary Figure 5). This indicates that the multiple interactions between the loops compensate for mutations that break a single interaction. It is interesting to note that the mutational robustness of these loops has been exploited in bioengineering applications, where insertion of an aptamer into one of the loops and randomization of the other allowed for the selection of synthetic riboswitches (Townshend et al., 2015). The identification of robust structural elements though high-throughput mutational data could be useful for identifying better targets for aptamer integration in other ribozymes.

### Epistasis plots are an informative approach to visualizing high-throughput activity data

Previous studies have reported comprehensive pairwise mutagenesis of ribozymes that provide interesting opportunities for comparison to the data presented here. For example, all pairwise mutations in a 42-nucleotide region of the same twister ribozyme were previously reported (Kobori and Yokobayashi, 2016). Compared to our experiments, these previous experiments used a later transcriptional time point (2h) and lower magnesium concentration (6mM). They did not calculate epistasis, and reported the Relative Activity of all double mutants using heatmaps similar to the figures presented here. The results were highly similar, and the authors were able to identify paired regions in the data. The similarity between the results illustrates the reliability of this sequencing-based approach, which is promising for future data sharing and meta-analysis efforts. In another prior work, all pairwise mutations in the glmS ribozyme were analyzed using a custom-built fluorescent RNA array (Andreasson et al., 2020). The power of this approach is that they were able to monitor self-cleavage over short and long time scales, which enables differentiating both very slow and very fast self-cleaving variants. While the authors did not calculate pairwise epistasis, they reported relative activity heatmaps and also “rescue effects” when the activity of a double mutant is sufficiently higher than the activity of a single mutant. This rescue analysis is very similar to positive epistasis, but only takes into account one mutation at a time. This analysis was also able to identify many of the know base-pair interactions and some tertiary contacts in the glmS ribozyme. In addition, they were able to observe some minor secondary structure rearrangement, where mutations in some nucleotides were able to rescue neighboring nucleotides by shifting the base-pairing slightly. The pairwise epistasis analysis presented here adds an additional approach to extract information from such high-throughput sequencing-based analysis of self-cleaving ribozymes. Unlike the rescue analysis, which can only identify positive interactions, the ability to detect negative epistatic interactions may help further identify structurally important regions for RNA sequence design and engineering efforts.

## CONCLUSION

We have determined the relative activity for all single and double mutants of five self-cleaving ribozymes and use this data to calculate epistasis for all possible pairs of nucleotides. The data was collected under identical co-transcriptional conditions, facilitating direct comparison of the data sets. The data revealed signatures of structural elements including paired regions and non-canonical structures. In addition, the comprehensiveness of the double mutants enabled identification of catalytic residues. Recently, there has been significant progress towards predicting RNA structures from sequence using machine learning approaches. The machine learning models are typically trained on structural biology data from x-ray crystallography, chemical probing (SHAPE), and natural sequence conservation. Self-cleaving ribozymes have been central to this effort. Our approach is similar to SHAPE in that it can be obtained with common lab equipment and commercially available reagents. The activity data presented provides information similar to natural sequence conservation, except that it provides quantitative effects of mutations, not just frequency. We hope that the activity-based data presented here will provide information not present in these other training data sets and help advance computational predictions.

## MATERIALS AND METHOD

### Mutational library design and preparation of self-cleaving ribozymes

Single-stranded DNA molecules used as templates for in vitro transcription were synthesized with 97% of the base of the reference sequence and 1% of the three other remaining bases at each position (Keck Oligo Synthesis Resource, Yale). The ssDNA library was made double stranded to allow for T7 transcription via low cycle PCR using Taq DNA polymerase.

### Co-transcriptional self-cleavage assay

The co-transcriptional self-cleavage reactions were carried out in triplicate by combining 20 μL 10X T7 transcription buffer (500 μL 1M Tris pH 7.5, 50 μL 1M DTT, 20 μL 1M Spermidine, 150 μL 1M MgCl2, 280 μL RNase Free water), 4 μL rNTP (25mM, NEB, Ipswich, Ma), 8 μL T7 RNA Polymerase-Plus enzyme mix (1,600 U, Invitrogen, Waltham, Ma), 160 μL nuclease free water, and 8 μL of double stranded DNA template (4 pmol, 0.5 μM PCR product) at 37°C for 30 minutes. The transcription and co-transcription self-cleavage reactions were quenched by adding 60 uL of 50 mM EDTA. The resulting RNA was purified and concentrated using Direct-zol RNA MicroPrep Kit with TRI-Reagent (Zymo Research, Irvine, Ca), and eluted in 7μL nuclease free water. Concentrations were determined via absorbance at 260 nm (ThermoFisher NanoDrop, Waltham, Ma), and normalized to 5μM. Reverse transcription reactions used 5 picomoles RNA and 20 picomoles of reverse transcription primer in a volume of 10 μL. RNA and primer were heated to 72 °C for 3 mins and cooled on ice. Reverse transcription was initiated by adding 4 μL SMARTScribe 5x First-Strand Buffer (TaKaRa, San Jose, Ca), 2 μL dNTP (10 mM), 2 μL DTT (20 mM), 2 μL phased template switching oligo mix (10 μM), and 2 μL SMARTScribe Reverse Transcriptase (200 units, TaKaRa) (Bendixsen et al., 2020). The mixture was incubated at 42 °C for 90 mins and the reaction was stopped by heating to 72 °C for 15 mins. The resulting cDNA was purified on a silica-based column (DCC-5, Zymo Research) and eluted into 7 μL water. Illumina adapter sequences and indexes were added using high-fidelity PCR. A unique index combination was assigned to each ribozyme and for each replicate. The PCR reaction contained 3 μL purified cDNA, 12.5 μL KAPA HiFi HotStart ReadyMix (2X, KAPA Biosystems, Wilmington, Ma), 2.5 μL forward, 2.5 μL reverse primer (Illumina Nextera Index Kit) and 5 μL water. Several cycles of PCR were examined using gel electrophoresis and a PCR cycle was chosen that was still in logarithmic amplification, prior to saturation. Each PCR cycle consisted of 98 °C for 10 s, 63 °C for 30 s and 72 °C for 30 s. PCR DNA was purified on silica-based columns (DCC-5, Zymo Research) and eluted in 22.5 μL water. The final product was then verified using gel electrophoresis.

### High-throughput sequencing

The indexed PCR products for all replicates were pooled together at equimolar concentrations based off of absorbance at 260 nm. Paired end sequencing reads were obtained for the pooled libraries using an Illumina HiSeq 4000 (Genomics and Cell Characterization Core Facility, University of Oregon).

### Sequencing data analysis

Paired-end sequencing reads were joined using FLASh, allowing ‘outies’ due to overlapping reads. The joined sequencing reads were analyzed using custom Julia scripts that implement a sequence-length sliding window to screen for double mutant variants of a reference ribozyme. Nucleotide identities for each mutant were identified and then counted as either cleaved or uncleaved based on the presence or absence of the 5’-cleavage product sequence. The relative activity (RA) was calculated as previously described (Kobori and Yokobayashi, 2016). Briefly, a fraction cleaved (FC) was calculated for each genotype in each replicate as FC=N_clv_/(N_clv_ + N_unclv_). This value was normalized to the reference/wild type fraction cleaved as RA = FC/FCwt. The RA values were averaged across the three replicates and then plotted as a heatmap. Epistasis interactions for each double mutant (i, j) were quantified as previously described (Bendixsen et al., 2017), where 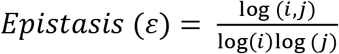. In order to eliminate false detection of epistasis interactions, values were filtered to eliminate instances where the difference between the double and any of the single mutants was less than 1-3σ of the overall distribution of differences between the single and double mutant relative activities. Values greater than 1 indicate positive epistasis, and values less than zero indicate negative epistasis. In order to eliminate false positive epistasis values where the.

### Correlation of thermodynamic stability of paired regions and observed mutational effects

Each base paired region was split into two separate RNA sequences containing only the nucleotides involved in base pairing, omitting nucleotides belonging to stem loops. Complex formation between each pair of strands at was analyzed in Nupack using Serrra and Turner RNA energy parameters in order to obtain minimum free energy values for each paired region (37°C, [1μM]). Using custom Julia scripts, the median relative activity for single mutations to each paired region was plotted as a function of the calculated free energy and a Pearson correlation coefficient was calculated.

## Acknowledgements

The authors acknowledge funding from the National Science Foundation (EH, grant number OIA-1738865, OIA-1826801), National Aeronautics and Space Administration (EH, grant number 80NSSC17K0738), and the Human Frontier Science Program (EH, Ref.-No: RGY0077/2019).

## Data Availability

Sequencing reads in FastQ format are available at ENA (PRJEB52899). Sequences, activity data, and computer code is available at GitLab (https://gitlab.com/bsu/biocompute-public/mut_12).

