## Supplementary Materials for "RNA sequence to structure analysis from comprehensive pairwise mutagenesis of multiple self-cleaving ribozymes"

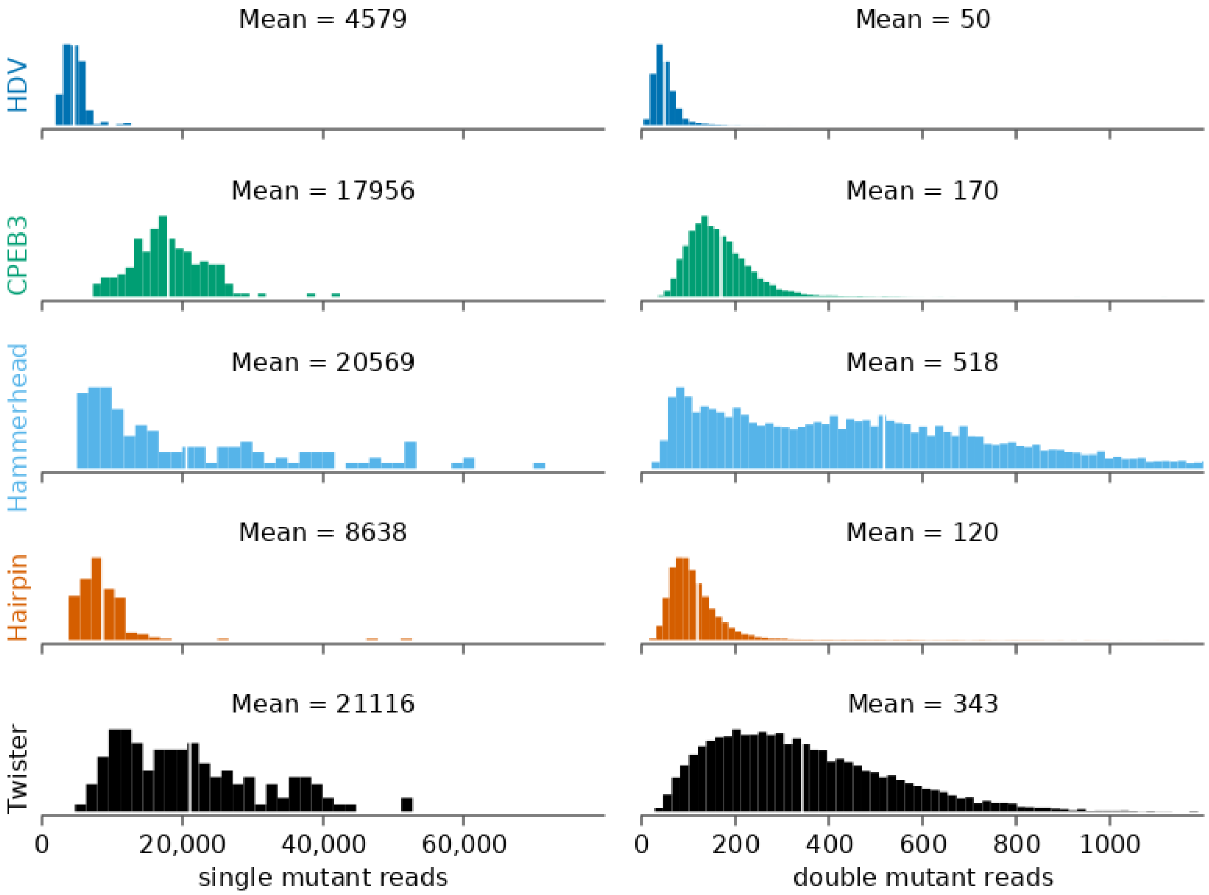


**Supplementary Figure 1:** Histogram of the distributions of read counts (read depth) for the single and double mutants matching to each ribozyme analyzed in this study (HDV, CPEB3, hammerhead, hairpin, twister).

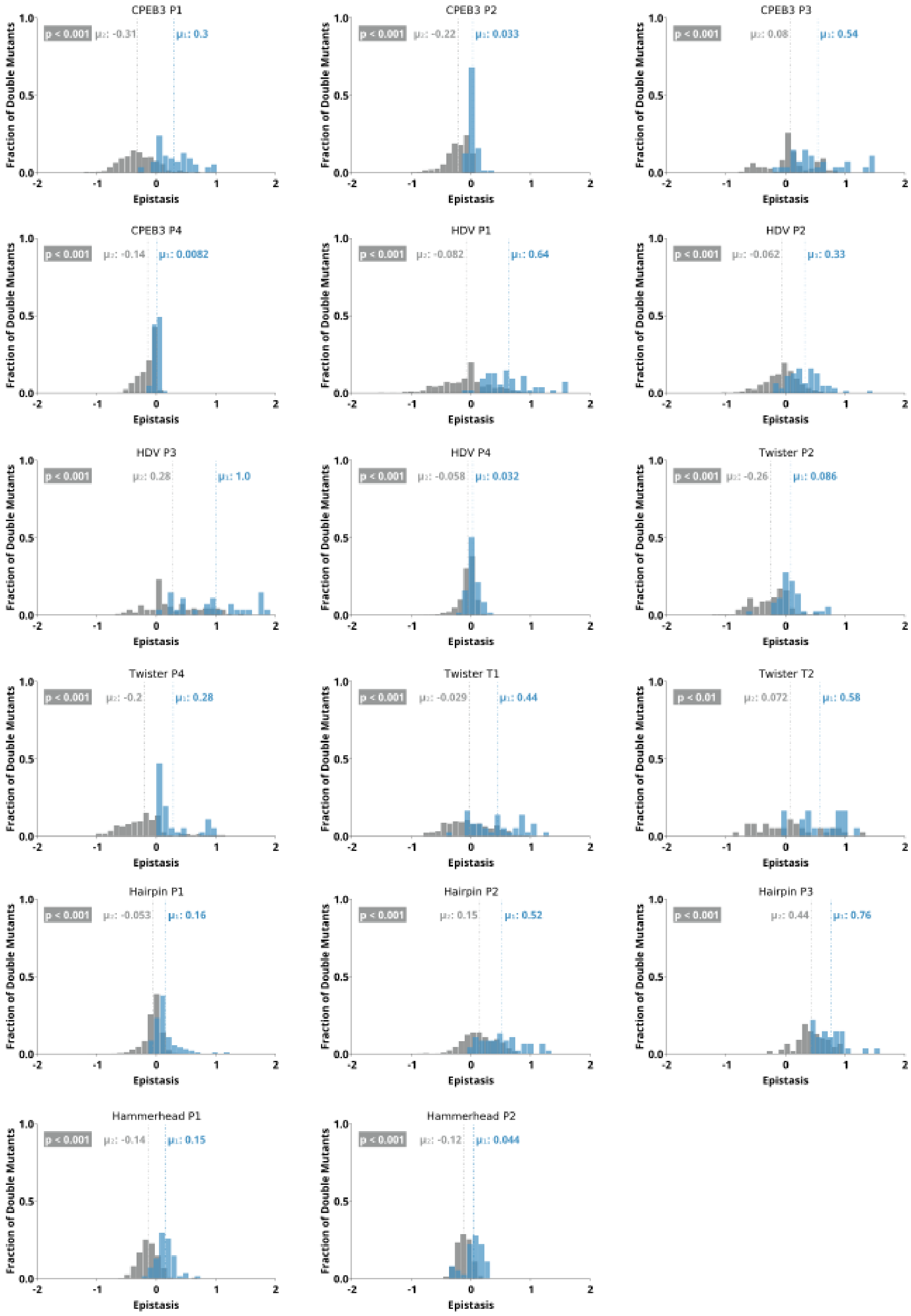


**Supplementary Figure 2:** Distributions for epistasis values seen on and off anti-diagonal in the epistasis heatmaps. The distributions of epistasis values along the anti-diagonal corresponding to double mutations between nucleotides involved in a Watson-Crick base-pair are shown in blue, and the epistasis values seen off diagonal are shown in gray.

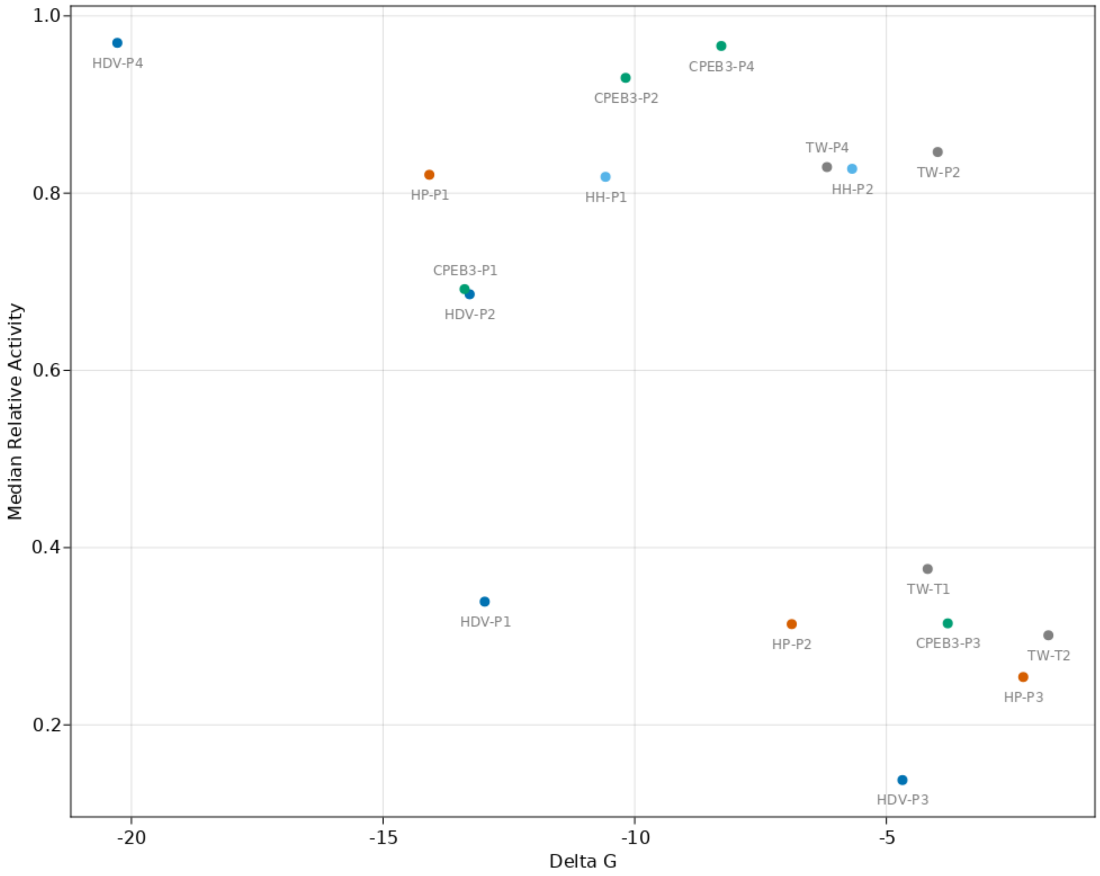


**Supplementary Figure 3:** Relationship between the Gibbs free energy (ΔG) of each base paired region belonging to the hairpin, hammerhead, CPEB3, HDV, and twister ribozymes, and the median relative activity of all single mutants within each base paired region (Pearson Correlation = -0.53).

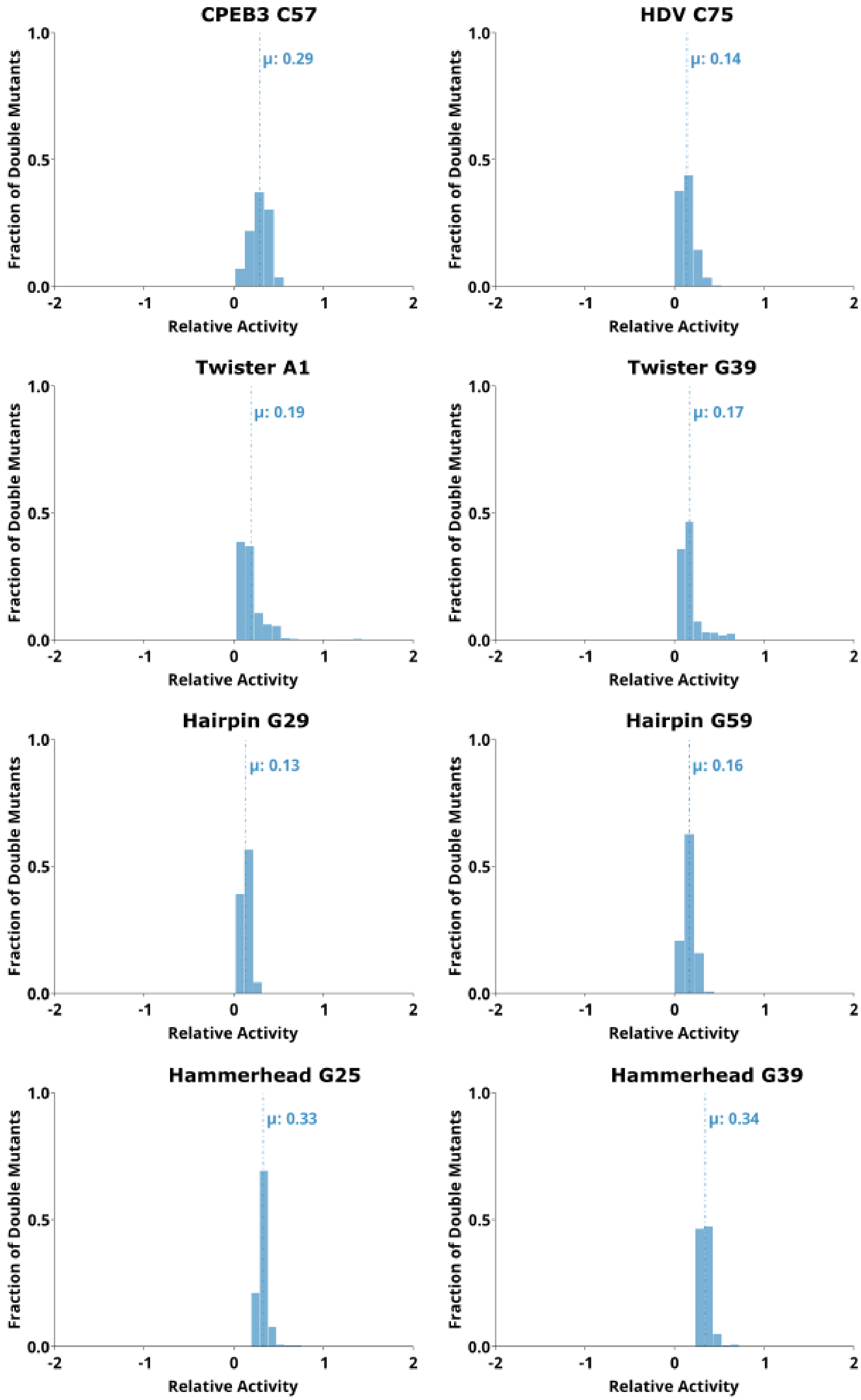


**Supplementary Figure 4:** Distributions of relative self-cleavage activity observed for sequences containing mutations to the catalytic nucleotides in the CPEB3, HDV, twister, hairpin, and hammerhead ribozymes.

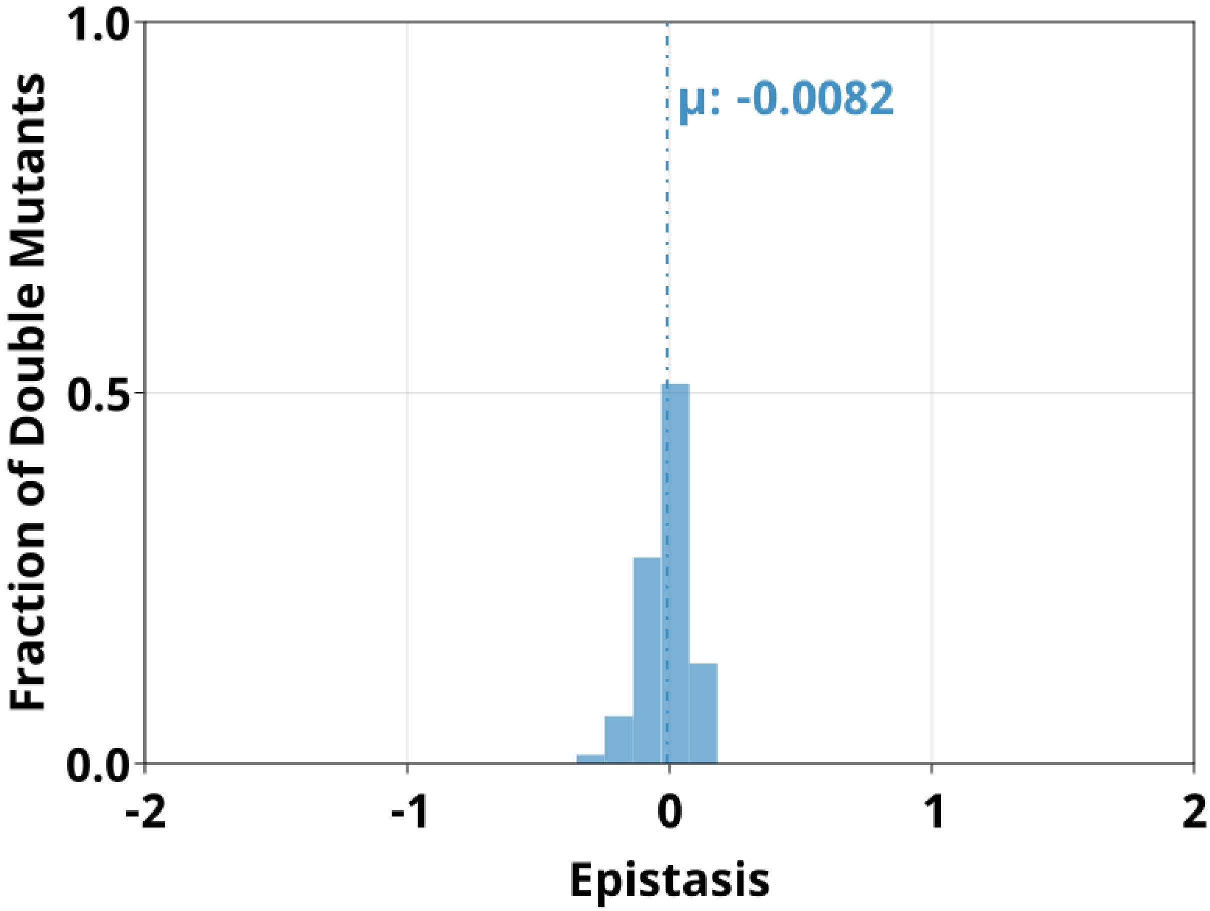


**Supplementary Figure 5:** Distribution of pairwise epistasis observed between the loops of P1 and P2 in the hammerhead ribozyme.
